# Computationally guided high-throughput design of self-assembling drug nanoparticles

**DOI:** 10.1101/786251

**Authors:** Daniel Reker, Yulia Rybakova, Ameya R. Kirtane, Ruonan Cao, Jee Won Yang, Natsuda Navamajiti, Apolonia Gardner, Rosanna M. Zhang, Tina Esfandiary, Johanna L’Heureux, Thomas von Erlach, Elena M. Smekalova, Dominique Leboeuf, Kaitlyn Hess, Aaron Lopes, Jaimie Rogner, Joy Collins, Siddartha M. Tamang, Keiko Ishida, Paul Chamberlain, DongSoo Yun, Abigail Lytoon-Jean, Christian K. Soule, Jaime H. Cheah, Alison M. Hayward, Robert Langer, Giovanni Traverso

## Abstract

Nanoformulations are transforming our capacity to effectively deliver and treat a myriad of conditions. However, many nanoformulation approaches still suffer from high production complexity and low drug loading. One potential solution relies on harnessing co-assembly of drugs and small molecular excipients to facilitate nanoparticle formation through solvent exchange without the need for chemical synthesis, generating nanoparticles with up to 95% drug loading. However, there is currently no understanding which of the millions of possible combinations of small molecules can result in the formation of these nanoparticles. Here we report the development of a high-throughput screening platform coupled to machine learning to enable the rapid evaluation of such nanoformulations. Our platform identified 101 novel self-assembling drug nanoparticles from 2.1 million pairings derived from 788 candidate drugs with one of 2686 excipients, spanning treatments for multiple diseases and often harnessing well-known food additives, vitamins, or approved drugs as carrier materials – with potential for accelerated approval and translation. Given their long-term stability and potential for clinical impact, we further characterize novel sorafenib-glycyrrhizin and terbinafine-taurocholic acid nanoparticles *ex vivo* and *in vivo*. We anticipate that this platform could accelerate the development of safer and more efficacious nanoformulations with high drug loadings for a wide range of therapeutics.

---

A large and increasing number of small molecular therapeutics suffer from poor water solubility,^1^ which can result in the formation of micron-sized colloidal aggregates^2,3^ that decrease bioavailability and therapeutic efficacy.^3,4^ The creation of solid drug nanoparticles through the addition of lipoid or polymeric stabilizers has provided a viable path to enable the delivery of such therapeutics,^5,6^ but this approach suffers from often complex metabolic profiles of the carriers as well as low drug loading, indicating that a majority of the administered material constitutes carrier rather than the desired active pharmaceutical ingredient.^7–9^ Recent work has revealed that co-self-assembly with azo and indocyanine dyes such as Congo red and IR783 can successfully stabilize a wide range of cancer drugs with high drug loading and thereby enhancing translational potential.^10,11^ We hypothesized that this nanoformulation technique is transferable to a wider range of drugs and excipients. Specifically, we postulated that drugs for various indications can be stabilized using a range of already FDA-approved excipients.^9,12^ This could potentially accelerate the translation of these novel systems through usage of carriers with well understood metabolism and safety. However, it is currently not understood which of the millions of possible combinations lead to formation of desired nanoparticles. To interrogate this question and test our hypothesis, we designed a novel high-throughput experimental platform coupled to state-of-the-art machine learning to identify pairs of drugs and excipients that will co-aggregate to form stable, self-assembled solid drug nanoparticles based on solvent exchange without the need for chemical synthesis (Figure 1A). Our system identified a total of 101 novel co-aggregated solid drug nanoparticles from 2.1 million possible pairings of 788 candidate drugs with one of 2686 excipients. We focused on using FDA-approved materials with potential applications ranging from cancer therapy to immunosuppression, asthma therapy, anti-viral, anti-malarial, and anti-fungal drug delivery. Recognizing the significant mortality and rapidly increasing incidence of hepatocellular carcinoma^13^ as well as the significant burden of onychomycosis^14^ with a paucity of novel therapeutic options, we focused further characterization on novel co-aggregated nanoparticles consisting of sorafenib-glycyrrhizin and terbinafine-taurocholic acid. Our *ex vivo* and *in vivo* proof-of-concept studies validate the potential of our platform to design safer and more efficacious nanoformulations based on approved carrier materials for a wide range of therapeutic applications with high drug loading and low production complexity.

**Figure 1:**
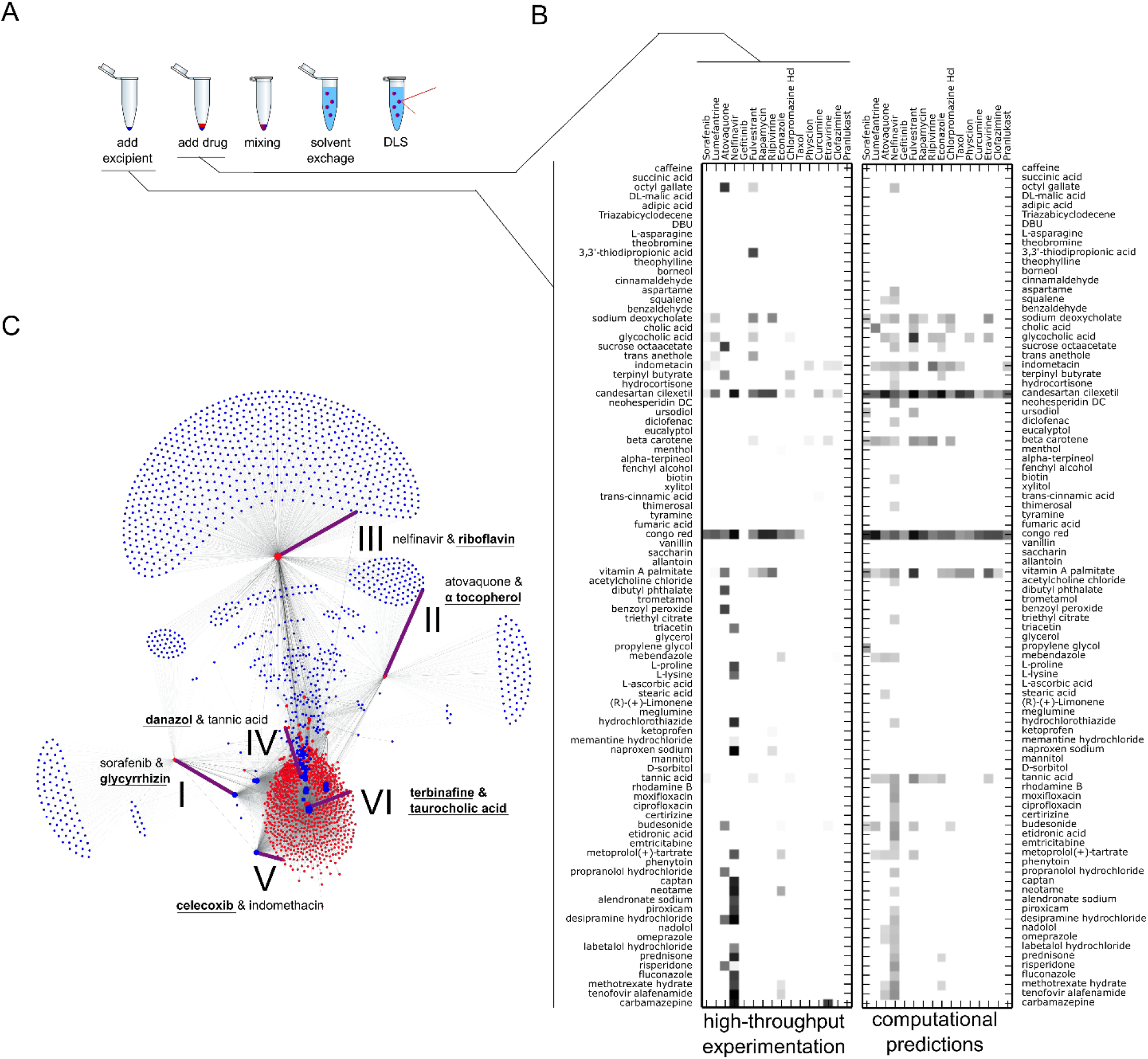
High-throughput creation of solid drug nanoparticles and computational prediction of drug-excipient pairs that lead to nanoparticle formation. **A** Schematic of experimental workflow to create nanoparticles using nanoprecipitation technique and rapid feasibility assessment using dynamic light scattering (DLS). **B** Left: High-throughput testing of all 1440 combinations of 16 drugs and 90 excipients (inactive ingredients, generally-recognized-as-safe food and drug additives, and other FDA-approved approved compounds). Color gradient indicates nanoparticle formation at physiological pH in PBS at room temperature, where white corresponds to no nanoparticle formation and shades of black correspond to relative size of formed nanoparticles. Right: Computational assessment of nanoparticle-forming potential of drug-excipient pairs according to their chemical structures, physicochemical properties, and pairwise interaction potential. Gradient indicates predictive confidence. We found good agreement between the computational assessments (right) and the real-world experimentation (left), with 91% of the experiments predicted correctly. **C** Using the here developed computational prediction model, we predicted 2.1 million pairs constituting all exhaustive combinations of 788 aggregating drugs each paired with one of 2686 excipients. The machine learning model predicts that a majority of the drugs and excipients will not interact and thereby result in colloidal aggregation and precipitation of the drugs. For 38,464 combinations (1.8%), the machine learning model predicted a potential interaction between the drug and the excipient that can lead to stabilized, self-assembled nanoparticle formation. Clear patterns are emerging, with some excipients (blue dots) enabling creation nanoparticles with many different drugs (red dots) while some other excipients can only selectively create nanoparticles with a single drug. Node size corresponds to the number of predicted excipients/drugs enabling formation of nanoparticles with this compound. Edge thickness corresponds to predictive confidence. Here further evaluated combinations are highlighted with purple edges, numbered (*cf*. Figure 2) and labelled with the corresponding drug and excipient constituting this pair. The novel component that was not previously used in the screen is highlighted in bold and underlined.

## High-throughput nanoparticle platform

Firstly, we needed to extract a set of candidate drugs that would be amenable to our formulation strategy by self-assembling into colloidal drug aggregates.^2,3^ A random forest machine learning model enabled us to rapidly predict the self-aggregation propensity of all small molecules in Drugbank 5^15^ with high specificity.^3,16^ A total of 788 drugs and other bioactive compounds were predicted to potentially self-aggregate in aqueous conditions and thereby represent candidate material for our nanoparticle formulation platform. We selected 20 compounds based on commercial availability and chemical diversity for our screening library. Four of these compounds did not show detectable aggregates on our platform and were discarded from further investigation (Supplementary Table 1). Secondly, a set of 90 excipients were selected according to chemical diversity, commercial availability, and biological safety from the FDA list of inactive ingredients,^12^ the FDA list of generally-recognized-as-safe ingredients,^17^ and currently approved drugs^15,18^ that were not predicted to be self-aggregating. This allowed us to generate and experimentally test 1440 novel formulations for their ability to form self-aggregating nanoparticles. We developed a high-throughput co-aggregation platform based on a liquid handling deck (Tecan Freedom Evo 150) coupled to a high-throughput dynamic light scattering platform (Wyatt Dyna Pro Plate Reader) to measure nanoparticle size. This system enabled us to rapidly screen 384-well plates of potential novel formulations using as little as 1 nanomole of drug or excipient for a single experiment replicate. After measuring all 1440 combinations of drugs and excipients in duplicates, 94 of those combinations (6.5%) showed nano-sized co-aggregate formation (Figure 1B). Interestingly, the data suggests a complex molecular recognition mechanism, with 21 excipients interacting only with a single drug each. Other excipients seem more accommodating and can form nanoparticles with a wide variety of drugs, for example the previously published^10^ positive control Congo red (10 drugs) and the newly identified drug candesartan cilexitel that acts as a stabilizer (9 drugs). From a drug-centric perspective, some therapeutics such as nelfinavir seem to be more prone to co-assembly (21 different nanoparticles formed) while none of our excipients was able to stabilize gefitinib or pranlukast.

## Machine learning for nanoparticle design

To extrapolate this dataset, we built a machine learning model based on a random forest classification model using chemical substructures and physicochemical properties of the drugs and excipients as well as autonomously interpreted, short molecular dynamic (MD) simulations to quantify non-covalent interaction potential. Conservative retrospective evaluation based on a “leave one drug out” policy^19^ had shown that a model based on all these three parameter types was most predictive (Supplementary Table 2) and that the random forest model architecture would consider all feature types relevant for prediction (Supplementary Figure 1). We confirmed that random forest models outperformed other, simpler machine learning models such as naïve Bayes and nearest neighbor (Supplementary Table 3). Furthermore, we were able to show that a y-shuffling derived adversarial straw model had no predictive power (Supplementary Table 4), indicating that our model identified meaningful chemical patterns in the dataset.^20^ Harnessing this machine learning model, we predicted the complete co-aggregation landscape of all 788 aggregating drugs combined with any of the 2686 available excipients, leading to 2.1 million formulations that we computationally assessed for their ability to form self-assembling nanoparticles. Given the infeasibility of running MD simulations for all these combinations, we applied a machine learning model trained exclusively on chemical and physicochemical properties with slightly lower retrospective performance but higher computational tractability (Supplementary Table 2). The machine learning model suggested a total of 38,464 combinations (1.8% of all possible pairs) to provide co-assembled nanoparticles. We observed a large heterogeneity in the interaction network, with many excipients (blue nodes) exclusively stabilizing a single drug (red nodes), while other excipients form central nodes that suggest a potential to stabilize multiple different drugs into effective nanoparticles. We set out to prospectively investigate some of these associations, aiming to cover different areas of the network and a wide range of different indications for the investigated drugs. We ultimately decided to test six of the predictions, specifically selecting high confidence predictions of novel excipients for drugs that we had already investigated (Figure2, I-III), new drugs for excipients that we had already investigated (Figure2, IV & V) and a combination of a novel drug and excipient (Figure2, VI).

**Figure 2:**
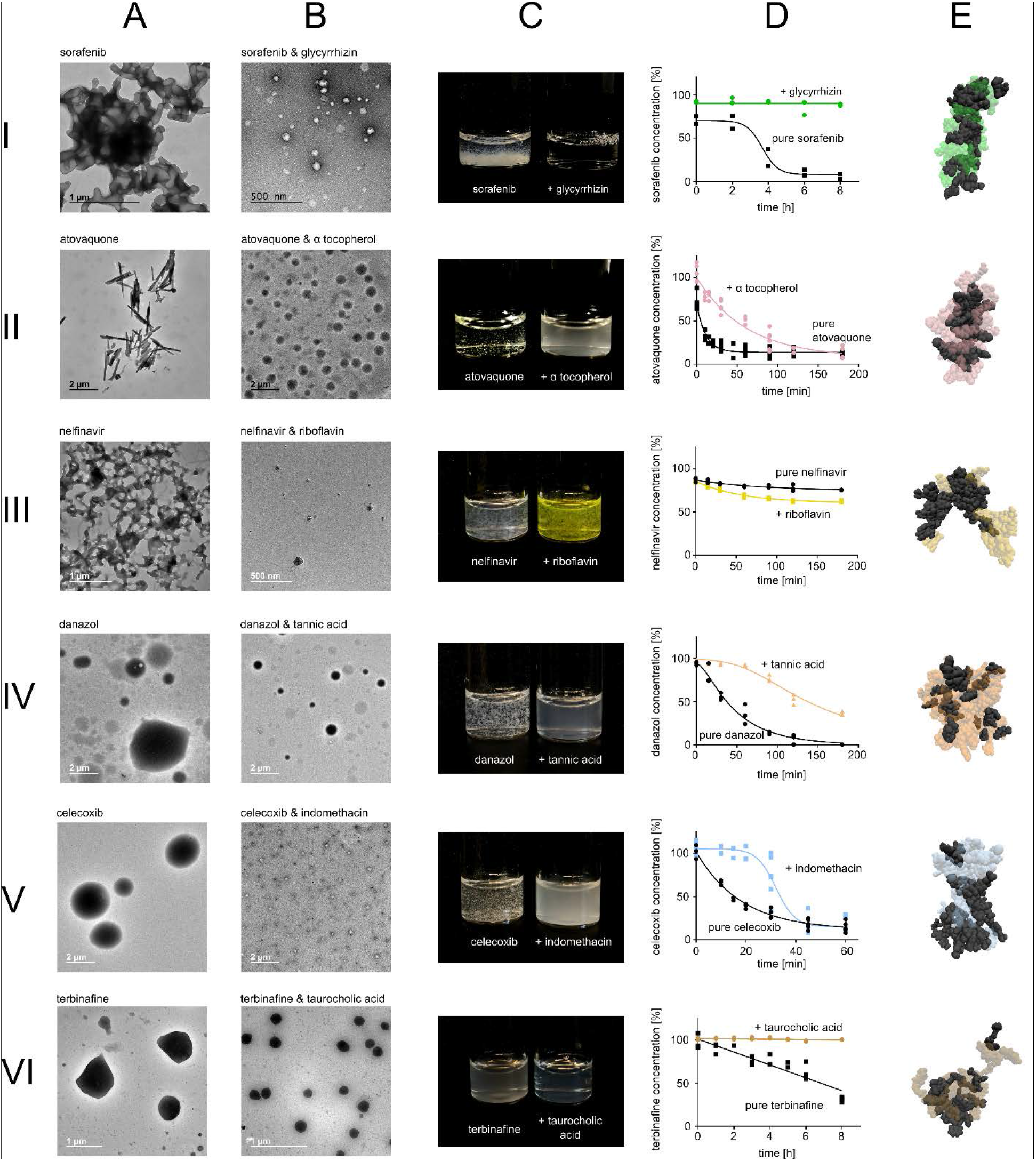
Novel and computationally predicted combinations of drugs and excipients evaluated for their ability to form solid drug nanoparticles. Numbering corresponds to edges highlighted in Figure 1C. Shown are TEM images of micron-sized aggregates formed by the pure drug (**A**) and TEM images of the nanoparticles formed by combining the drugs and excipients (**B**). Photos show dispersion of the nanoparticles compared to unformulated drug during concentration escalation experiments (**C**). Precipitation dynamics of drugs alone or in nanoparticle formulations, as measured using HPLC. (**D**). Short molecular dynamic simulations map non-covalent interaction potential between drugs and excipients (**E**). In the MD visualizations, drugs are visualized through black Van der Waals-spheres, while excipients are visualized through transparent and colored Van der Waals-spheres.

## *In silico* designed nanoparticles improve dispersion

We next characterized the novel predicted pairs, which consisted of novel excipients (I-III), novel drugs (IV and V) or a combination of a novel excipient coupled to a novel drug (VI). First, we measured dynamic light scattering (DLS) of all six drugs alone and found that they indeed formed large micron-sized, polydisperse aggregates in PBS containing 1% DMSO (Supplementary Table 5) that grew rapidly and eventually precipitated out of solution. These colloidal aggregations were further confirmed using transmission electron microscopy (TEM), revealing a wide range of complex microstructures formed by these medications in aqueous environments (Figure 2A). All the computationally suggested excipients for co-aggregation reduced particle size by at least two-fold (Supplementary Table 5). TEM confirmed that co-aggregation formed homogeneous populations of self-assembled nanoparticles (Figure 2B). To further characterize these co-aggregates, we escalated concentrations of drug and particles at equal rates to enable visual assessment of the nanoparticle stability and improved dispersion (Figure 2C). With the exception of nelfinavir-riboflavin, all our novel nanoparticular co-aggregates show clear or milky dispersions at high drug concentrations up to 1 mM, while the unformulated drugs precipitated rapidly. To analyze the kinetics of precipitation and long term stability of the nanoparticles, we sampled drug concentration of nanoparticles and unformulated drug with HPLC measurements at fixed concentrations (Figure 2D). The six drugs displayed different precipitation dynamics, with four exhibiting an exponential loss of nano-dispersed drug. Sorafenib seems to be more stable until it suddenly precipitates. Terbinafine shows the highest variation among different samples but overall precipitates in a more linear fashion. The novel nanoparticles exhibit dramatically altered precipitation dynamics and significantly slow the precipitation or even fully stabilize the drug dispersion (Figure 2D). We further investigated this behavior with more extensive MD simulations to study the ability of the excipients to halt aggregation of the drugs *in silico* through non-covalent interactions (Figure 2E). In agreement with the experimental data, all excipients were able to reduce the aggregation propensity of the drugs with the exception of riboflavin for nelfinavir. We selected sorafenib-glycyrrhizin and terbinafine-taurocholic acid nanoparticles for further characterization and *ex vivo* and *in vivo* applications given the superb long-term stability of these two co-aggregate systems (Figure 2D) as well as the clinical relevance of using innovative delivery approaches for these drugs in the treatment of hepatocellular carcinoma^13^ and onychomycosis.^14^

## Terbinafine particles for antifungal applications

Onychomycosis is the most common nail disorder observed in clinical practice.^14^ This common fungal infection is caused by dermatophytes, nondermatophytes and yeast. We hypothesized that stabilized nanoformulations of an approved therapeutic could enable topical applications with improved tissue penetration to maximize local efficacy. Terbinafine is a first-line treatment of cutaneous mycoses, but its effectiveness is reduced in topical applications due to decreased tissue penetration.^21^ We hypothesized that our novel terbinafine-taurocholic acid particles could retain therapeutic effects of terbinafine while improving skin penetration. In-depth TEM assessment revealed distinct aggregation dynamics with populations of different types of low density nanoparticles forming. Three different types of nanostructures emerge that dominate different phases of the nanoparticle formation process, as revealed by timed TEM imaging (Figure 3A). We used ultracentrifugation for purification of the particles and noticed that the particles exhibited low density and precipitated in the supernatant. Analytics of this precipitate revealed high encapsulation efficiency (80 ± 1.9%) and even higher drug loading (96 ± 1.6%) (Figure 3B). Scanning Transmission Electron Microscopy-Energy Dispersive Spectroscopy (STEM-EDS) showed that the particles contained substantial amounts of chlorine from the excipient taurocholic acid – providing further evidence that the nanoparticles are indeed co-aggregates of drugs and excipients (Supplementary Figure 2). We next tested whether the nanoparticles would retain the fungistatic effect of terbinafine against *Candida albicans*, a major source of onychomycosis in immunocompromised individuals,^22^ and found that our particles halted biofilm formation with comparable efficiency compared to free terbinafine (Figure 3C and D). Interestingly, our particles improved the skin permeation of terbinafine in Franz Diffusion cell experiments by two-fold, supporting the potential of our novel particles to improve local enhanced bioavailability and amplify topical delivery of terbinafine (Figure 3E).

**Figure 3:**
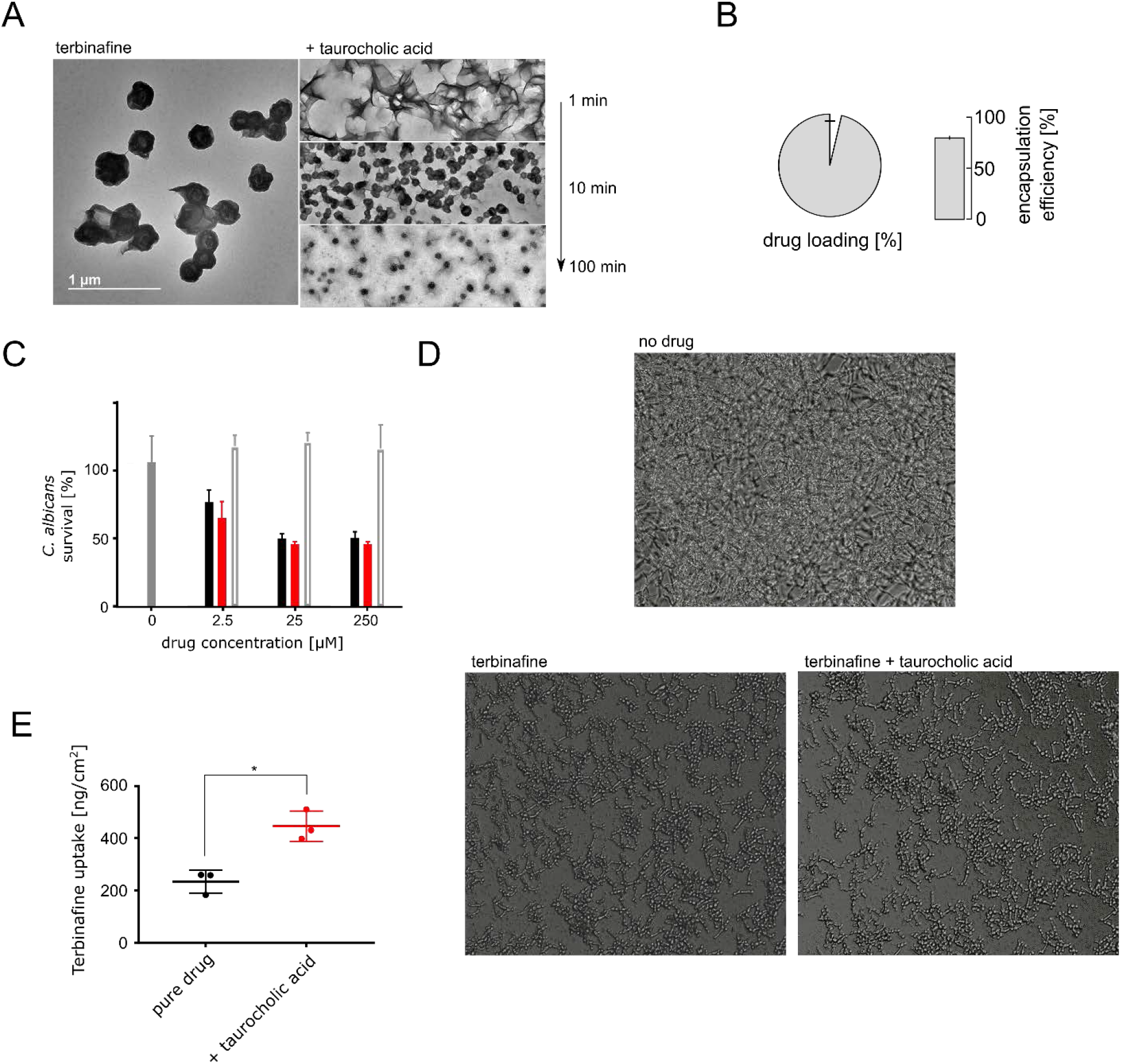
Characterization and evaluation of terbinafine & taurocholic acid particles. **A** TEM images of terbinafine alone and nanoparticles formulated together with taurocholic acid. TEM images taken at different timepoints after self-assembly initiation are depicted on the right. Scale bar corresponds to 1 μm, scale consistent across images. **B** Ultracentrifugation and LC/MS analytics of pellets enabled determining high drug loading (93%) and encapsulation efficiency (80%) of terbinafine & taurocholic acid nanoparticles. **C** *C. albicans* survival after treatment with terbinafine alone (positive control, black bars), terbinafine & taurocholic acid nanoparticles (red bars), pure taurocholic acid (formulation control, light gray bars), 1% DMSO in PBS (untreated control, gray bars), n = 5. **D** Microscopy images of *C. albicans* after 17h of nanoparticle treatment or terbinafine treatment compared to untreated control. **E** Skin uptake of terbinafine into porcine flank skin in Franz Diffusion cell measurements, n = 3, *p* = 0.028 two-sided T-test.

## Sorafenib nanoparticles improve anti-cancer efficacy

In a second proof-of-concept, we aimed at translating our novel sorafenib nanoformulations. Sorafenib is a multi-kinase inhibitor used as a targeted therapy of several types of cancer. Significantly, sorafenib is the only approved first-line treatment for advanced hepatocellular carcinoma (HCC).^23^ There is a significant clinical need for new treatments for HCC given the increasing incidence^13^ and poor prognosis.^23^ Nanoparticles have previously shown early promising results at improving sorafenib efficacy,^11^ which motivated us to use our platform to identify novel sorafenib nanoparticles. We compared our predicted stabilizer glycyrrhizin with three nanoparticle-forming excipients identified during the screening (candesartan cilexitel, indomethacin, tannic acid) as well as low confidence prediction meloxicam, chosen from a rational perspective as it represents a COX inhibitor like the hit indomethacin. These excipients were chosen given their well understood safety or potentially previous association with sorafenib therapy. Glycyrrhizin, although a known 11β-hydroxysteroid dehydrogenase inhibitor associated with reversible hypermineralocorticoid-like effects after intensive consumption, is considered a safe food and drug ingredient with daily consumption of up to 3.6 mg/kg.^24^ Sorafenib has been previously associated with development of hypertension^25^ and has been clinically applied together with candesartan to counter this effect.^26,27^ Similarly, cyclooxygenase (COX) inhibitors have been studied as synergistic enhancers of anticancer effects of sorafenib,^28,29^ in particular for meloxicam.^30^ Therefore, formulating sorafenib with such drugs might constitute an important step towards excipient-free formulations.^5,31^ Sorafenib constitutes a prime example for such a development given that additional therapeutic agents are likely to be necessary to boost the therapeutic efficiency and decrease the susceptibility to drug resistance development for this treatment.^30^ All investigated excipients created self-assembled nanoparticles with radii below 100nm (Figure 4A, Supplementary Table 6). All particles were stable and improved drug dispersion compared to unformulated drug (Figure 4B), with all but meloxicam fully-stabilizing the dispersion over the investigated time span. The nanoparticles also exhibited significantly improved cytotoxicity in human liver carcinoma HUH7 cells over a wide range of concentrations compared to unformulated sorafenib (Figure 4C). With the exception of tannic acid, we were able to refute that any of the used excipients show significant cytotoxicity in HUH7 alone (Supplementary Figure 3) and thereby ruled out additive effects leading to increased cytotoxicity. Instead, we hypothesized that the nanoparticles augment drug uptake compared to the larger micro-scale structures of sorafenib alone (Supplementary Figure 4).^4^ We investigated our tannic acid, indomethacin and glycyrrhizin nanoparticles for their ability to impact sorafenib uptake. While all particles seemed to increase uptake, the glycyrrhizin particles most significantly impacted drug availability and almost doubled cytosolic sorafenib content (Figure 4D). We confirmed that sorafenib-glycyrrhizin particles did not inhibit sorafenib from engaging with one of its main targets Raf1 by observing an equivalent inhibition of MEK phosphorylation compared to free sorafenib (Supplementary Figure 5). We were able to further purify our particles using ultracentrifugation, which revealed that the nanoparticles had very high drug loading (95 ± 1.3%) and equally notable encapsulation efficiency (93 ± 2.1%) (Figure 4E). STEM-EDS showed that the particles contained substantial amounts of fluorine from sorafenib, revealing that the particles are enriched with drug as expected from the analytics (Supplementary Figure 6). DLS showed that the particles were stable in serum and medium and were not disrupted by shearing forces when administered through a 29 Gauge needle, or ultracentrifugation followed by ultrasound-mediated re-dispersion (Supplementary Figure 7).

**Figure 4:**
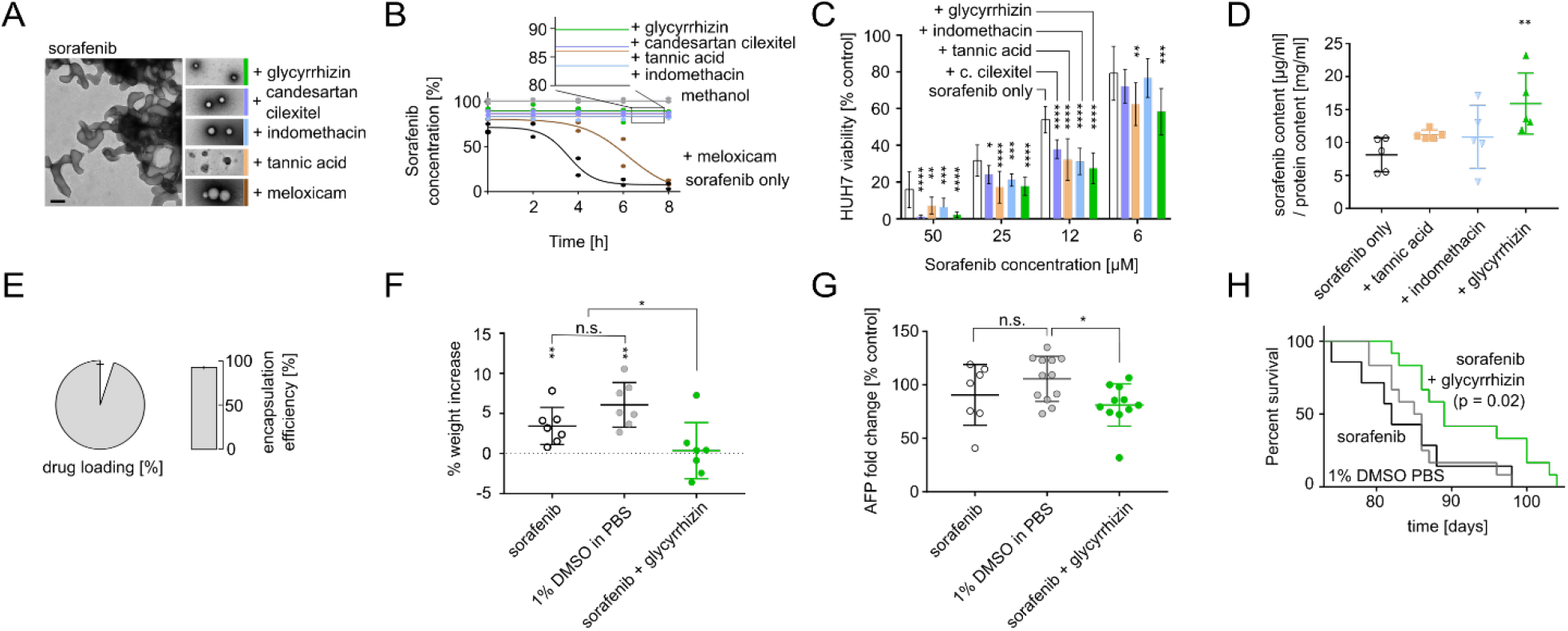
Characterization and *in vivo* application of sorafenib nanoparticles. **A** TEM images of colloidal aggregates formed by unformulated sorafenib compared to TEM images of nanoparticles formulated with glycyrrhizin, candesartan cilexitel, indomethacin, tannic acid, and meloxicam. Black scale bar corresponds to 200 nm, scale consistent over all images. **B** Dispersion of sorafenib alone and in nanoformulation with excipients as measured by HPLC over time. Percentages for nanoparticles reflect encapsulation efficiency. **C** HUH7 human hepatocellular carcinoma cell survival after 48 h of treatment with different nanoparticles or free sorafenib over a wide range of drug concentrations. n = 4, two-tailed T test. *p* = 0.0013 two-way ANOVA. D Nanoparticles improve cell membrane permeation of sorafenib in HUH7 cells after 2h incubation, most significantly for glycyrrhizin (p = 0.006 two-tailed T-test), n=5, *p* = 0.016 one-way ANOVA. E Ultracentrifugation of nanoparticles and HPLC analytics revealed high drug loading (93%) and encapsulation efficiency (91%) for sorafenib particles with glycyrrhizin. F Mice treated with sorafenib-glycyrrhizin particles had no significant tumor growth as indicated by no increase in bodyweight after four weeks of treatment (*p* = 0.79, one-sample T test) while mice treated with oral sorafenib and the 1% DMSO in PBS control showed significantly increased tumor burden (*p* < 0.007, one-sample T-test), *p* = 0.0063 one-way ANOVA, *p* = 0.0032 Dunnett’s multiple comparisons test, n = 7. G Sorafenib-glycyrrhizin treated mice showed significantly reduced AFP levels after 4 weeks of treatment compared to 1% DMSO in PBS control treatment (*p* = 0.027, Dunnett’s multiple comparisons test) while oral sorafenib did not reduce AFP levels (*p* = 0.29, Dunnett’s multiple comparisons test), *p* = 0.045 one-way ANOVA, n ≥ 7. H Kaplan-Meyer analysis shows that mice treated with sorafenib-glycyrrhizin nanoparticles show longer morbidity-free survival (*p* = 0.02, log-rank Mantel-Cox test) compared to oral sorafenib and 1% DMSO in PBS control treatment groups (*p* = 0.83, log-rank Mantel-Cox test), n ≥ 7.

We refined our formulation protocol to consistently generate 3 mg / mL dispersions of sorafenib-glycyrrhizin nanoparticles (Supplementary Figure 8) that enabled us to dose 30 mg / kg for IV injection.^23,32^ In toxicity analysis, our formulation did not induce abnormal liver function after three injections of elevated dosages of 60 mg/kg (Supplementary Table 7). Next, we set out to compare the *in vivo* antitumor efficacy of our sorafenib-glycyrrhizin nanoparticles compared to sorafenib alone in an established genetic mouse model of spontaneous HCC,^23,32^ the most common type of liver cancer. The cancer is induced by an overexpression of human ΔN90-β-catenin and human MET genes in the mouse liver and represents an aggressive model of HCC characterized by rapid formation of a large number of tumors.^32^ HCC bearing mice were treated three times a week with 1% DMSO in PBS, oral sorafenib, or sorafenib-glycyrrhizin. After four weeks of treatment, no tumor growth was observed in mice treated with our sorafenib-glycyrrhizin nanoparticles as indicated by no increase in bodyweight (Figure 4F). All control groups showed significant tumor burden and increase in bodyweight (Figure 4F and Supplementary Figure 9). In line with the weight gain data, mice treated with sorafenib-glycyrrhizin nanoparticles had significantly lower alpha-fetoprotein (AFP) levels as a marker of transformed hepatocytes compared to controls (Figure 4G and Supplementary Figure 10). Calculation of survival fractions using the Kaplan-Meier method revealed that mice treated with sorafenib-glycyrrhizin nanoparticles had significantly longer morbidity-free survival (p = 0.02, log-rank Mantel-Cox test) compared to PBS-treated mice or mice receiving only sorafenib (Figure 4H). We also demonstrated that glycyrrhizin alone or the route of administration did not explain this increase in efficacy (Supplementary Figure 11). Taken together, our results demonstrate improved efficacy of the computationally designed nanoparticles and serve as proof-of-concept for our platform to enable alternative modes of delivery for life-saving therapeutics.

## Conclusion

Nanoparticles are one of the most promising technologies to enable the delivery of therapeutics with otherwise challenging LADMET profiles.^33^ However, the translation of nanoparticles can be hindered by high production complexity and low drug loading. Recent progress in utilizing co-assembly with small molecular dyes to form solid drug nanoparticles from chemotherapeutics has provided nanoparticles with high drug loading and ease of production through simple solvent exchange.^10,11^ Unfortunately, only a limited number of stabilizing excipients have been reported and these focused investigations do not enable to anticipate which of the millions of possible combinations of drugs with small molecular excipients would lead to nanoparticle formation. With our platform, we broaden the scope of this technology for a wide array of different types of drugs for treating various indications as well as harnessing well-understood and FDA-approved excipients such as vitamins, nutrients, food compounds, and drugs. We have characterized in-depth two examples in which these nanoparticle formulations can potentially enable a novel mode of delivery with potentially improved efficacy and reduced toxicity.

An important challenge and opportunity lies in the context-sensitive nature of the self-assembly process. Buffer conditions such as pH, temperature, and salt concentrations might significantly impact the aggregation and co-aggregation propensity of the here described materials.^2,3,34^ We here focused on physiological pH at room temperature, but variations of these conditions for different drugs and excipients needs to be considered during manufacturing and could broaden the scope for materials that are not currently accessible through our platform. Considering aggregation context can provide further opportunities for the development of adaptive systems that aggregate specifically *in situ*, for example in gastric conditions.^35^ This might also ultimately enable responsive systems: for example, we had observed that some of our aggregates grow when heated (Supplementary Figure 12), potentially due to Ostwald ripening and elevated collision rates.^36^ If such conditions can be controlled, dynamic changes to the nanoparticles might be harnessed for adaptive delivery solutions.^37^

Future research will include development of methods to enable the rapid prediction of important physical and biological properties of these novel nanoparticles. Investigating focused sets of nanoparticles can enable the prediction of expected sizes of the formed particles.^11^ We have provided further evidence here that nanoparticle size can correlate with particle stability (Figures 2 and 4). Furthermore, our data suggests predictive uncertainty measures (*cf*. meloxicam in Figure 4) as well as MD assessments (*cf*. riboflavin in Figure 2) being able to further delineate nanoparticle stability. If possible, the prediction or modification of material characteristics such as anticipated biodistribution, immunogenicity, and surface properties will dramatically accelerate the translation of these novel nanoparticle systems and provide a viable option for the delivery of life-saving therapeutics with unfavorable LADMET profiles.

We believe that such small molecule-based nanoparticle systems are a fruitful addition to the thriving nanomedicine armamentarium, enabling rapid generation of highly loaded nanocarriers for life-saving therapeutics with otherwise unfavorable LADMET profiles.^10,11^ This platform provides a rapid pipeline to identify feasible combinations of such drugs with safe inactive ingredients or other FDA approved materials with improved understanding of their biological implications and potential for rapid translation.^9^ Taken together, with the ability to choose specific stabilizers for an extremely wide range of drugs, we expect that the approach reported herein and extensions thereof will be an important step towards personalized drug delivery.^9^

## Methods & Materials

### Identification of candidate drugs

A random forest machine learning model was trained on publicly available aggregation data provided by the Brian Shoichet laboratory (bkslab.org) as previously described.^38^ Briefly, all structures were encoded through Morgan fingerprints (radius 3, 2048 bits) and physicochemical descriptors (RDKit). A random forest classifier (scikit-learn, 1000 trees, no maximum features) was training and evaluated using ten-fold cross validation.^3^ To predict the self-aggregation behavior of approved drugs, all 1857 small molecular structures in the category “approved” were extracted in SMILES format from DrugBank 5.0.^15^ To enable aggregation prediction, these were also encoded through Morgan fingerprints (radius 3, 2048 bits) and physicochemical descriptors (RDKit). Missing values in the physicochemical descriptions were estimated through mean values using a scikit-learn imputation module. Aggregation likelihood was estimated through the number of trees in the random forest model that consider a DrugBank compound to be an aggregator (maximum predictive confidence).

### High-throughput formulation assessment

10mM stock solutions of aggregating drugs and excipients were created in sterile DMSO and stored at −20 °C. For every experiment, 1 μl of the drug stock solution was mixed with 1 μl of the stock solution of any of the excipients (1:1) in a 96 well plate using a Tecan Freedom Evo 150 liquid handling deck. Mixing of the two droplets was ensured through brief shaking and centrifugation. Subsequently, we rapidly added 198 μl of sterile-filtered and degassed PBS for solvent exchange and mixed the novel formulation through repeated pipetting. Replicate samples of 75 μl were subsequently transferred into a 384 well plate for high-throughput dynamic light scattering assessment on a Wyatt Dyna Pro Plate Reader at 25 °C using 5 independent acquisitions of 5 s duration. Data was processed by calculating the median size observed for a specific drug and requiring at least a two-fold size reduction to indicate co-aggregation. Measures of polydispersity and replicate variance were considered but did not lead to improved predictive performance.

### Machine learning

Chemical structures for all drugs in SMILES representation were extracted from DrugBank 5.0.^15^ Data for excipients was extracted from the FDA according to previously published protocols.^9^ Compounds were described according to radial chemical substructures (Morgan Fingerprint, radius 4, 1024 bits)^39^ and calculated physicochemical properties (RDKit).^40^ Concatenating the substructure description and the physicochemical properties for every drug and every excipient generates a 2440-dimensional description of a drug-excipient formulation. Additionally, short MD simulations were run and automatically analyzed to assess the enthalpic, non-covalent interaction potential between drugs and excipients. In brief, atomic partial charges for drugs and excipients were derived from antechamber and parmchk2 and the net charge was calculated using the OpenEye Quacpac AM1-BCC method. Two excipient and two drug molecules were randomly positioned using packmol and amber’s tleap module. After energy minimization of the system, short 20ns simulations were run in OpenMM 7.2.1 in OBC2 implicit solvent with non-periodic boundary conditions and no cutoff distance. A Langevin Integrator was used at 300K with friction coefficient of 1/ps. The timestep of the simulation was 2 fs and the trajectory was saved every 10 ps, creating 2,000 frames. These frames were processed with MDTraj to calculate heavy atom distances between drugs and excipients and derive summary statistics (maximum, minimum, average, as well as maximum first derivative) for each pair. Furthermore, potential and kinetic energy of the last frame were calculated and an interaction potential based on pharmacophore typing was derived on the last frame. Overall, this gave us an additional 20 parameters to characterize drug and excipient pairs as input to the machine learning platform. These 2460dimensional numerical characterizations of a formulation served as input for the random forest classification machine learning model (scikit-learn) with 500 trees and using ⌊√2460⌋ = 49 features per classification tree.

### Transmission electron microscopy

Nanoparticles were created at 50 μM drug concentration in 1% DMSO PBS. Samples were observed using TEM with or without negative staining. In either case, 7 μl of nanoparticle solution was transferred onto a 200 mesh copper grid coated with a continuous carbon film, and excess solution was wicked away after 60 seconds. In case of negative staining, 10 μl of staining solution (phosphotungstic acid 1% in water) was added and excess solution was wicked away immediately. Subsequently another round of staining was performed using the same volume of staining solution and excess solution was wicked away after 40 seconds from the edges of the samples. Finally, all samples were then dried at room temperature before imaging. Afterwards, the grid was mounted on a JEOL single tilt holder. Imaging was done on a JEOL 2100 FEG microscope. The microscope was operated at 200 kV and with a magnification in the range of 3,000 to 60,000 for assessing particle shape and size and atomic arrangement. All images were recorded on a Gatan 2kx2k UltraScan CCD camera. STEM imaging was done using a HAADF (high-angle annular dark field) detector with 0.5 nm probe size and 12 cm camera length. A X-Max 80mm^2^ EDX (Oxford Instrument, UK) was used to determine the chemical information of the samples.

### Concentration escalation experiments

Unformulated drug and nanoparticles were created by mixing 5 μl of drug stock solution in DMSO with 5 μl of excipient stock solution in DMSO or pure DMSO and performing solvent exchange by adding 990 μl sterile filtered and degassed PBS. These experiments were repeated at escalating concentrations until either 5mM concentrations were reached or drug alone visibly precipitated. Images were taken at 1 mM for atovaquone, 500 μM for celecoxib, 5 mM for terbinafine, 1 mM for sorafenib, 500 μM for danazol and 500 μM for nelfinavir.

### Analytics

Dispersion studies were conducted with 1 ml of 250 μM formulations in glass tubes, except for altered concentrations for celecoxib (500 μM) or nelfinavir (50 μM) to enable studying dynamics at the same time scales compared to other drugs in spite of slower/faster aggregation dynamics. Formulations were sampled at pre-determined timepoints and the sample was diluted in acetonitrile or methanol to ensure full solubility of the drug and disruption of the nanoparticles for further analytics. Sorafenib (250μM), nelfinavir (50 μM), and celecoxib (500 μM) formulations were analyzed using an Agilent 4.6 x 50 mm EC C-18 Poroshell column with 2.7 μm particles, maintained at 50 °C. The optimized mobile phase consisted of 0.1 % formic acid in water (A) and acetonitrile (B) using a flow rate of 1 mL/min. Gradient separation was achieved over a 13 minute run time at the following parameters: 0 min 95% A and 5% B; 4.4 min 50% A and 50% B; 6.4 min 50% A and 50% B; 9 min 20% A and 80% B; 9.5 min 20% A and 80% B; and 10 min 95% A and 5% B. The injection volume was 10 μl, and the selected ultraviolet (UV) detection wavelength was 248 nm with no reference parameters at an acquisition rate of 40 Hz. Atovaquone formulations (250 μM) were analyzed using a Phenomenex Sphereclone 4.6 x 250 mm ODS (I) column with 5 μm particles, maintained at 50 °C. The optimized mobile phase consisted of 5% water, 15% methanol, and 80%acetonitrile using a flow rate of 1 mL/min over a 15 minute run time. The injection volume was 10 μl, and the selected ultraviolet (UV) detection wavelength was 295 nm with no reference parameters at an acquisition rate of 40 Hz. Danazol (250 μM) formulations were analyzed using an Agilent Zorbax Eclipse XDB C-18 4.6 x 150 mm column with 5 μm particles, maintained at 35 °C. The optimized mobile phase consisted of 15%, 0.1% formic acid in water and 85% methanol using a flow rate of 1 mL/min over a 6 minute run time. The injection volume was 5 μl, and the selected ultraviolet (UV) detection wavelength was 285 nm with no reference parameters at an acquisition rate of 10 Hz. Terbinafine (250 μM) formulations were analyzed via HPLC using an Agilent Zorbax Eclipse XDB C-18 4.6 x 150 mm column with 5 μm particles, maintained at 40 °C. The optimized mobile phase consisted of 0.1 % formic acid in water (A) and methanol (B) using a flow rate of 1 mL/min. Gradient separation was achieved over a 10 minute run time (3 min post run) at the following parameters: 0 min 70% A and 30% B; 5 min 30% A and 70% B. The injection volume was 10 μl, and the selected ultraviolet (UV) detection wavelength was 242 nm with no reference parameters at an acquisition rate of 10 Hz.

### Additional molecular dynamics simulations

For selected combinations of drugs and excipients, we converted SMILES compound representations to Tripos mol2 files using Openbabel with B3LYP/6-31G* partial charge calculation with GAFF force field and systematic rotor search for generating 3D conformations. Missing parameters were checked with the parmchk2 module in Ambertools. Using packmol, five random system configurations were generated for 20 molecules of each pair (20+20=40 molecules total), or 20 molecules of drug only. A topology and coordinate file was generated with the tleap module in Ambertools. OpenMM 7.3.1 was used to create a 20ns simulation under OBC2 implicit solvent at 1 bar, 300K, with periodic cutoff distance of 1nm after initial minimization.

### *Candida albicans* XTT assay and microscopy

*Candida albicans* were incubated in sterile 50 g/L Difco YPD broth overnight at 30 °C on an orbital shaker at 300 rpm. Centrifuging at 3220 g for 5 mins was used to extract the fungus. Washing was performed twice with PBS. The final pellets were re-suspended in 20mL RPMI1640 (Sigma Aldrich). The fungi concentration was determined using a hemocytometer with bright-field microscopy at 40x magnification. Fungi were seeded at concentration of 1 million per well in 96-well plates. Terbinafine-taurocholic acid particles or free terbinafine were added at drug concentrations of 250 μM, 25 μM and 2.5 μM were added to different wells in quadruplicates. 1% DMSO PBS was used as buffer control and 70% isopropanol was treated as positive control. Plates were wrapped with parafilm and incubated at 37 °C. Fungi viability was evaluated using an XTT assay after 24h of incubation. To this end, XTT was prepared as a saturated solution at 0.5 g/L in sterile PBS under light protection and subsequently sterile filtered with a 0.22-mm pore size filter. XTT solution was aliquoted in 10 ml Falcon tubes and stored at −80 °C. Before a measurement, 1 μL of a 10 mM menadione solution in acetone was added to the XTT solution and used to substitute the media. Absorption was measured at 490nm to determine background signal per well. Plates were subsequently incubated for another 2h at 37 °C under light protection and signal was measured at 490nm absorption to determine fungus viability. For microscopy, 40x bright-field images were taken after 17h incubation using 25 μM free terbinafine, 25 μM terbinafine-taurocholic acid nanoparticles or 1% DMSO PBS buffer control at 37 °C.

### Skin uptake of terbinafine particles

All *ex vivo* studies were approved by the Massachusetts Institute of Technology Committee on Animal Care. Fresh porcine ear samples were provided by the Massachusetts Institute of Technology facilities and were washed three times with PBS and mounted on a Franz Diffusion cell (FDC-400 flat flange, 15 mm orifice diameter, mounted on an FDC diffusion drive console providing synchronous stirring at 350rpm, Crown Glass Co. Inc., Sommerville, N.J., USA) by gluing the epidermis side of the skin to the donor chamber and subsequent mounting onto the receiver chamber that was preloaded with PBS. Terbinafine-taurocholic acid nanoparticle or free terbinafine were added at 250 μM in 1% DMSO PBS and loaded to the donor chambers in triplicates. 1% DMSO in PBS served as buffer control. Parafilm was used to prevent evaporation from both the receiver and the donor chamber. After 4h of incubation, donor solutions were removed and washing with PBS was used to remove excess drug from the skin. The perfusion area was cut into squares and weighed and measured for further analysis. Skin samples were loaded into 2 mL homogenizing tubes (Fisher Scientific, Fisherbrand™ Pre-Filled Bead Mill Tubes) with methanol at a 1:2 ratio and homogenized at 6000 rpm for 40s x 3. Homogenate was analyzed for drug content using Ultra-Performance Liquid Chromatography-Tandem Mass Spectrometry (UPLC-MS/MS). Analysis was performed on a Waters ACQUITY UPLC®-I-Class System aligned with a Waters Xevo® TQ-S mass spectrometer (Waters Corporation, Milford MA). Liquid chromatographic separation was performed on an Acquity UPLC® BEH C18 (50mm × 2.1mm, 1.7 μm particle size) column at 50 °C. The mobile phase consisted of aqueous 0.1% formic acid, 10mM ammonium formate solution (Mobile Phase A) and acetonitrile: 10 mM ammonium formate, 0.1% formic acid solution (95:5 v/v) (Mobile Phase B). The mobile phase had a continuous flow rate of 0.6 mL/min using a time and solvent gradient composition. The initial composition, 95% Mobile Phase A, was held for 0.25 minutes. Afterwards, the composition was changed linearly to 5% Mobile Phase A and 95% Mobile Phase B until 1.00 minutes. The composition was held constant at 95% Mobile Phase B until 2.75 minutes. At 3.00 minutes the composition returned to 95% Mobile Phase A, where it remained for column equilibration for the duration of the run, ending at 4.00 minutes. The mass to charge transitions (m/z) used to quantitate terbinafine were 292.294>141.167 and 292.294>93.181 for quantitation and confirmation respectively. For internal standard, naftitine, 288.26>117.17 and 288.26>141.17 m/z transitions were used for quantitation and confirmation respectively. Sample introduction and ionization was by electrospray ionization (ESI) in the positive ionization mode. Waters MassLynx 4.1 software was used for data acquisition and analysis. Stock solutions were prepared in methanol at a concentration of 500 μg/mL. A twelve-point calibration curve was prepared in analyte-free, blank skin homogenate ranging from 1.25-5000 ng/mL. 100 μl of each sample was spiked with 200 μl of 250 ng/mL internal standard in acetonitrile to elicit protein precipitation. Samples were vortexed, sonicated for 10 minutes, and centrifuged for 10 minutes at 13,000 rpm. 200 μl of supernatant was pipetted into a 96-well plate containing 200 μl of water.

### HUH7 cell survival assessment

HUH7 cells were plated at 10,000 cells in 96 well plates and incubated overnight to allow adhesion. Cell medium was changed and cells were treated with various concentrations of nanoparticles for 48h. These concentration series were generated by creating a dilution series of the DMSO stock solutions and then performing nanoparticle generation at altered stock concentrations. We performed two independent experiments with n=4 replicates. Cell viability was measured through quantitation of ATP using CellTiter-Glo (Promega).^41^

### MEK phosphorylation quantification

Infrared secondary antibodies, IRdye 680RD goat anti-mouse IgG and IRdye 800CW anti-rabbit (both 1:1000) were purchased from LiCor. Phospho-MEK1/2 (Ser217/221) Antibody (9121) and MEK1/2 Antibody (9122) were purchased from Cell Signaling Technology. For In-Cell Western, cellular proteins were quantitated *in situ* based on infrared intensity. Samples were immunolabelled with an infrared conjugated IgG secondary antibody using standard immunofluorescence protocol. Briefly, cells were fixed with 4 % (v/v) formalin in dH2O (Sigma) for 15 minutes at room temperature, washed with PBS, permeabilized with 0.25 % (v/v) TritonX-100/PBS for 2 minutes, washed with PBS and then blocked with 4 % (w/v) bovine serum albumin in PBS. Primary and secondary antibodies were incubated in blocking buffer for 1 h at room temperature. Samples were subsequently imaged using an Odyssey Fc Infrared Imaging System (LiCor). The resulting signal intensity was subsequently quantified by using the Odyssey CLx Image Studio Analysis Software.

### Sorafenib uptake experiments

HUH-7 cells were plated in 12-well plates at 20,000 cells (1ml) and incubated for 24h to allow adhesion. Cell medium was changed (900 μl) and cells were treated with (100 μl, 10x) 100μM sorafenib, 100μM of novel nanoparticle formulations, or buffer control (0.1% DMSO PBS) and incubated for 2h. Afterwards the supernatant was removed and cells were carefully washed 3 times using a 2.5% Tween-80 PBS solution, which was shown previously to dissolve sorafenib aggregates while not resulting in changes of cell adhesion or morphology according to microscopy images. Afterwards, cells were lysed with Pierce RIPA Buffer (ThermoFisher Scientific) and transferred. Subsequently, lysate was lyophilized and re-dissolved in methanol (200 μl). The solution was centrifuged (10min at 3200RPM) and supernatant was subsequently analyzed using HPLC for sorafenib content. AUC values were converted into sorafenib content (μg / ml) according to a standard curve and normalized according to protein content per replicate (mg / ml) from an orthogonal BCA assay.

### *In vivo* cancer study

All *in vivo* studies were approved by the Massachusetts Institute of Technology Committee on Animal Care. Female FVB/N mice were purchased from Charles River Laboratories. Animals were maintained in a conventional barrier facility with a climate-controlled environment on a 12-h light/12-h dark cycle, fed ad libitum with regular rodent chow. Hepatocellular carcinoma was induced as previously described.^23,32^ Briefly, plasmids encoding human ΔN90-β-catenin, human MET, and SB transposase were hydrodynamically injected intravenously into 6–7-week-old mice. Plasmids were kindly provided by Dr. Xin Chen (UCSF, San Francisco, CA). Plasmids were amplified and isolated in endotoxin free conditions (< 5 EU/mg) by Aldevron (Fargo, ND). 5 weeks after injections serum levels of alpha-fetoprotein (AFP) were analyzed as HCC marker and animals were stratified in several groups. Treatment was started six weeks after HCC induction, and each group of mice was treated three times a week for four weeks with one of the following treatments: intravenous injections of 30 mg/kg sorafenib-glycyrrhizin nanoparticles, sorafenib suspensions at 30mg/kg in PBS with 1% DMSO delivered by oral gavage, intravenous injections of buffer control (1% DMSO in PBS), intravenous injections of 30 mg/kg sorafenib solubilized in a 50:50 ethanol-cremophor mixture, or intravenous injections of glycyrrhizin in 1% DMSO in PBS as a vehicle control. Mice were evaluated twice daily for signs of adverse effects related to tumor burden, change in bodyweight, and other clinical signs of discomfort such as labored breathing, lethargy, lack of appetite, cachexia, diarrhea, poor grooming, hunched appearance and lack of nest building. After the four week treatment period, AFP levels and bodyweight changes were recorded to evaluate treatment success.

### AFP quantification

Mouse blood was collected into BD Vacutainer® EDTA tubes, incubated on ice for 10 min and then centrifuged at 2000g for 20 min at 4°C. 3 μl of serum (the supernatant) was mixed with 47 μl PBS (Gibco) and 10 μl 6X reducing SDS buffer (Boston Bio Products), the mixture was heated at 68°C for 30 min, and 10 μl of the mixture were subjected to SDS/PAGE. Proteins were transferred to Nitrocellulose membranes (BioRad) and blocked with Li-COR Odyssey® Blocking Buffer (TBS) for 1 h at RT. Biotinylated anti-mouse AFP antibody (R&D Systems, BAF5369) was reconstituted at 0.2 mg/mL in sterile PBS and used at 1:2000 dilution. The binding was done for either 2 h at RT or overnight at 4°C. Protein bands were visualized by incubation with secondary antibodies labelled with infrared fluorophores (Li-COR IRDye® 680LT Streptavidin, 926-68031, dilution 1:10,000) for 30 min at RT. Membranes were scanned on a Li-COR Odyssey Scanner and fluorescence intensity was subsequently quantified using the ImageJ software. Band density was normalized to background per gel. The normalized band density for each sample was divided by the average normalized band density for no treatment controls to assess relative band density, reflecting the difference in AFP expression levels in the treated mice vs control mice. The relative change of this relative band density level before and after treatment was used to assess treatment success.

### Toxicity assessment

Two healthy, 6 week old, female FVB/N mice were treated with elevated dosages of 60 mg/kg sorafenib-glycyrrhizin nanoparticles for one week (three administrations). Blood was sampled after the final administration for serum chemistry assessment. We screened seven markers associated with liver damage (alanine aminotransferase ALT, aspartate aminotransferase AST, alkaline phosphatase ALP, albumin, gamma-glutamyl transferase GGT, total protein, direct bilirubin) by the Massachusetts Institute of Technology Division of Comparative Medicine.

## ACKNOWLEDGMENTS

D.R. is a Swiss National Science Foundation Fellow (Grants P2EZP3_168827 and P300P2_177833). Y.R. is grateful to the MIT Skoltech Initiative for financial support. A.R.K. is grateful to the PhRMA foundation postdoctoral fellowship for financial support. We thank the Koch Institute Swanson Biotechnology Center for technical support, specifically the High Throughput Sciences Facility, Nanotechnology Materials Facility, the Animal Imaging and Preclinical Testing core, and the Histology core. This work was supported in part by the Koch Institute Support (core) Grant P30-CA14051 from the National Cancer Institute and by the NIH Grant EB000244. We are grateful to OpenEye for providing us with an OpenEye Academic License. We are grateful to Hormoz Mazdiyasni for technical support and to Miguel Jimenez for providing access to C. albicans and helpful discussions throughout the study.

## CONTRIBUTIONS

D.R., R.L. and G.T. conceived the study. D.R., Y.R., A.R.K., R.L, and G.T. designed experiments. D.R. and J.W.Y. performed *in silico* experiments. D.R., R.C., J.W.Y., N.N., R.M.Z., T.E., J.L.H. performed *in vitro* experiments. D.R., R.C., A.G. performed *in vivo* experiments. Y.R., A.R.K., T.v.E., A.L.-J., C.K.S., J.H.C supported *in vitro* experiments. E.M.S, D.L, J.C., S.M.T, K.I., P.C. and A.M.H. supported in vivo experiments. D.S.Y. performed TEM imaging and K.H, A.L., J.R. performed pharmaceutical analytics. D.R., R.L. and G.T. wrote the manuscript with contributions from the other authors. All authors approved the final version of this manuscript.

## DATA AVAILABILITY

The authors declare that the data supporting the findings of this study are available within the paper and its supplementary information files.

## CODE AVAILABILITY

All code for this study will be made available upon request to the corresponding author.

