## Supplementary Material for "Computationally guided high-throughput design of self-assembling drug nanoparticles"

[1] Koch Institute for Integrative Cancer Research, Massachusetts Institute of Technology, Cambridge, MA 02139 (USA)
[2] Division of Gastroenterology, Department of Medicine, Brigham and Women's Hospital, Harvard Medical School, Boston, MA 02115 (USA)
[3] Division of Engineering Science, University of Toronto, Toronto, ON M5S 2E4 (CAN)
[4] Robert Frederick Smith School of Chemical and Biomolecular Engineering, Cornell University, Ithaca, NY 14853 (USA)
[5] Biomedical Engineering Program, Faculty of Engineering, Chulalongkorn University, Bangkok, 10330 (THA)
[6] Department of Electrical Engineering & Computer Science, Massachusetts Institute of Technology, Cambridge, MA 02139 (USA)
[7] Skolkovo Institute of Science and Technology, Moscow, 121205 (RUS)
[8] Department of Mechanical Engineering, Massachusetts Institute of Technology, Cambridge, MA 02139 (USA)

**Supplementary Table 1:** Drugs that failed to show aggregation and were therefore not amenable to our formulation approach.

| Drug | DLS radius [nm] |
|---|---|
| dyphylline | 65 |
| simvastatin | 80 |
| budenoside | 60 |
| minoxidil | 70 |

**Supplementary Table 2:** Retrospective machine learning performance using 10x "leave one drug out" evaluation of various tested descriptors for classification. The full model relying on chemical descriptors (FP) and molecular dynamics (MD) performs significantly better than any sub-model, although the "only

FP” model (using radial chemical substructure descriptors and physicochemical descriptors but no MD simulations) is competitive with a similar retrospective performance. *p* corresponds to two-sample student T-test of the ten repeats comparing the full model with the respective model in the row.

| | MCC | | F1 | | Precision | | Accuracy | |
|---|---|---|---|---|---|---|---|---|
| | **mean** | ***p*** | **mean** | ***p*** | **mean** | ***p*** | **mean** | ***p*** |
| **FP & MD** | 0.25 | -- | 0.30 | -- | 0.28 | -- | 0.90 | -- |
| **only FP** | 0.23 | 0.001 | 0.29 | 0.002 | 0.27 | 0.03 | 0.89 | 4e-5 |
| **only MD** | 0.08 | 3e-18 | 0.13 | 2e-18 | 0.14 | 1e-17 | 0.88 | 4e-11 |

**Supplementary Table 3:** Retrospective machine learning performance using 10x “leave one drug out” evaluation of various tested models for classification. Random forest models outperform simpler models such as Naïve Bayes and kNN. *p* corresponds to two-sample student T-test of the ten repeats comparing the model with the respective control model in the row.

| | MCC | F1 | Precision | Accuracy |
|---|---|---|---|---|
| **Random Forest** | 0.25 | 0.30 | 0.28 | 0.90 |
| **Naïve Bayes** | -0.03 | 0.03 | 0.03 | 0.87 |
| **kNN** | 0.13 | 0.20 | 0.14 | 0.82 |

**Supplementary Table 4:** Adversarial control model with y shuffling behaves randomly and does not predict self-aggregation, which suggests that the here developed machine learning model identifies molecular patterns associated with co-aggregation. *p* corresponds to two-sample student T-test of the ten repeats comparing the model with the adversarial control model through y shuffling.

| | MCC | | F1 | | Precision | |
|---|---|---|---|---|---|---|
| | **mean** | ***p*** | **mean** | ***p*** | **mean** | ***p*** |
| **Model** | 0.25 | -- | 0.30 | -- | 0.28 | -- |
| **Adversial control** | -0.01 | 1e-18 | 0.03 | 1e-19 | 0.04 | 1e-16 |

**Supplementary Table 5:** DLS results of computationally predicted self-assembling nanoparticles.

| Drug | Sample | Radius [nm] | Reduction factor |
|---|---|---|---|
| **Sorafenib** | drug alone | 846.0 | 18.7 |
| | + glycyrrhizin | 45.3 | |
| **Atovaquone** | drug alone | 910.4 | 8.9 |
| | + a tocopherol | 102.4 | |
| **Nelfinavir** | drug alone | 1102.0 | 9.4 |
| | + riboflavin | 117.5 | |
| **Danazol** | drug alone | 351.2 | 5.9 |
| | + tannic acid | 59.4 | |
| **Celecoxib** | drug alone | 646.1 | 11.1 |

| | | | |
|---|---|---|---|
| | + indomethacin | 58.1 | |
| **Terbinafine** | drug alone | 596.3 | 4.4 |
| | + taurocholic acid | 134.2 | |

**Supplementary Table 6:** DLS results of further characterized sorafenib nanoparticles.

| **Drug** | **Sample** | **Radius [nm]** | **Reduction factor** |
|---|---|---|---|
| **Sorafenib** | drug alone | 920.0 | **-----** |
| | + glycyrrhizin | 42.1 | **21.8** |
| | + candesartan cilexitel | 38.1 | **24.1** |
| | + tannic acid | 56.2 | **16.3** |
| | + indomethacin | 65.2 | **14.11** |
| | + meloxicam | 99.1 | **9.28** |

**Supplementary Table 7:** Serum chemistry assessment of liver toxicity associated markers in two healthy female 6 week old FVB/N mice. Mice received three injections of elevated dosages of 60 mg/kg sorafenib-glycyrrhizin particles within one week and showed no adverse symptoms.

| **Maker** | **Mouse 1** | **Mouse 2** | **Reference** |
|---|---|---|---|
| **ALT [IU/L]** | 42.0 | 45.0 | 17.0-77.0 |
| **AST [IU/L]** | 46.0 | 54.0 | 54.0-298.0 |
| **ALP [IU/L]** | 96.0 | 84.0 | 35.0-96.0 |
| **Albumin [g/dL]** | 3.0 | 2.7 | 2.5-4.8 |
| **GGT [IU/L]** | 0.0 | 0.0 | -- |
| **Total protein [g/dL]** | 5.2 | 5.0 | 3.5-7.2 |
| **Direct bilirubin [mg/dL]** | 0.0 | 0.0 | -- |

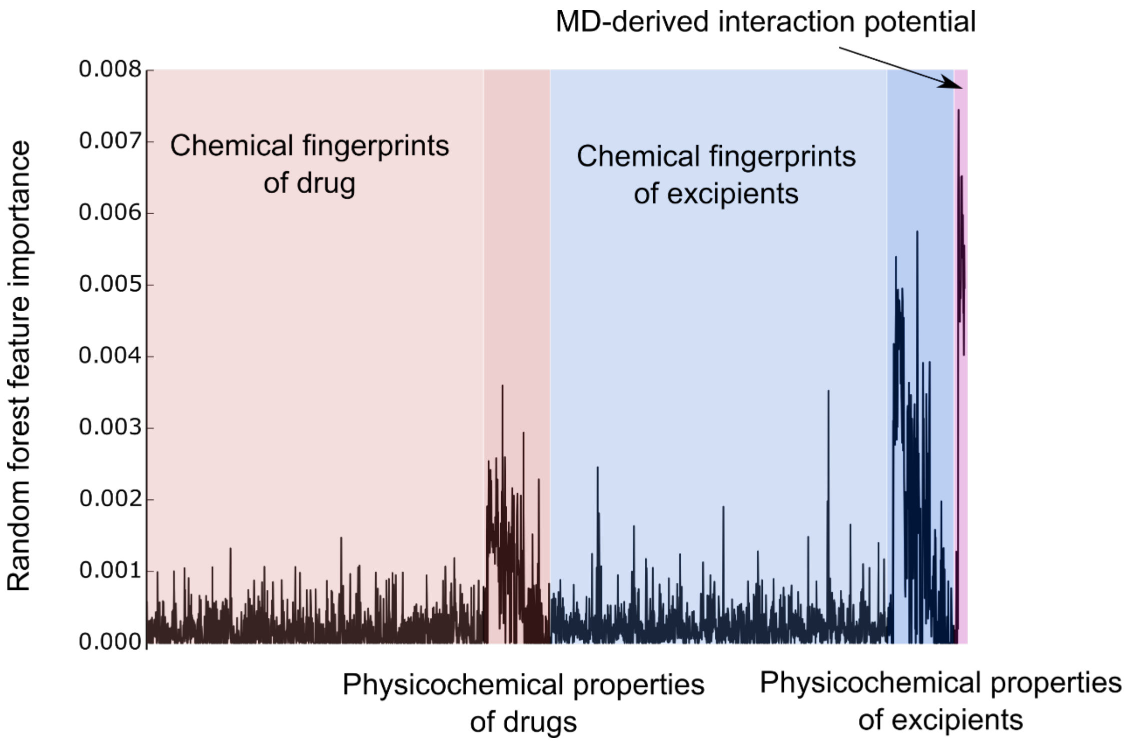


**Supplementary Figure 1:** Random forest feature importance of the final model for the 2460 numerical descriptors of the novel formulations. As can be seen here, the model considers both chemical and physicochemical properties of both the drug and the excipient as well as the MD-derived interaction potentials.

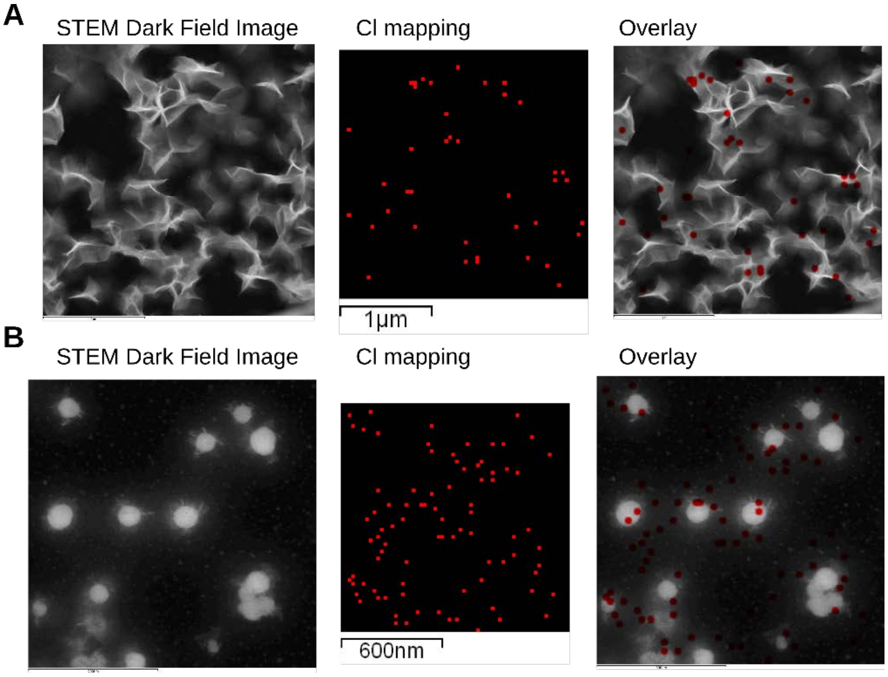

**Supplementary Figure 2:** Scanning Transmission Electron Microscopy-Energy Dispersive Spectroscopy (STEM-EDS) shows that terbinafine-taurocholic acid nanoparticles are rich in chlorine from taurocholic acid, both in cases of early aggregates (**A**) as well as for late-stage aggregates **(B).**

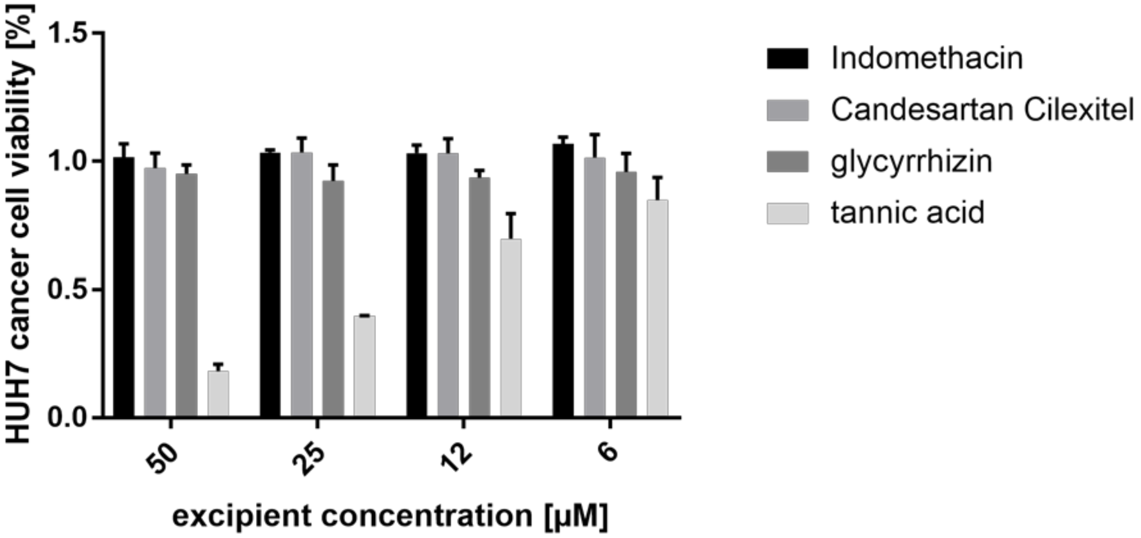


**Supplementary Figure 3:** HUH7 survival after 48 h of treatment with nanoparticle-forming excipients alone at different concentrations.

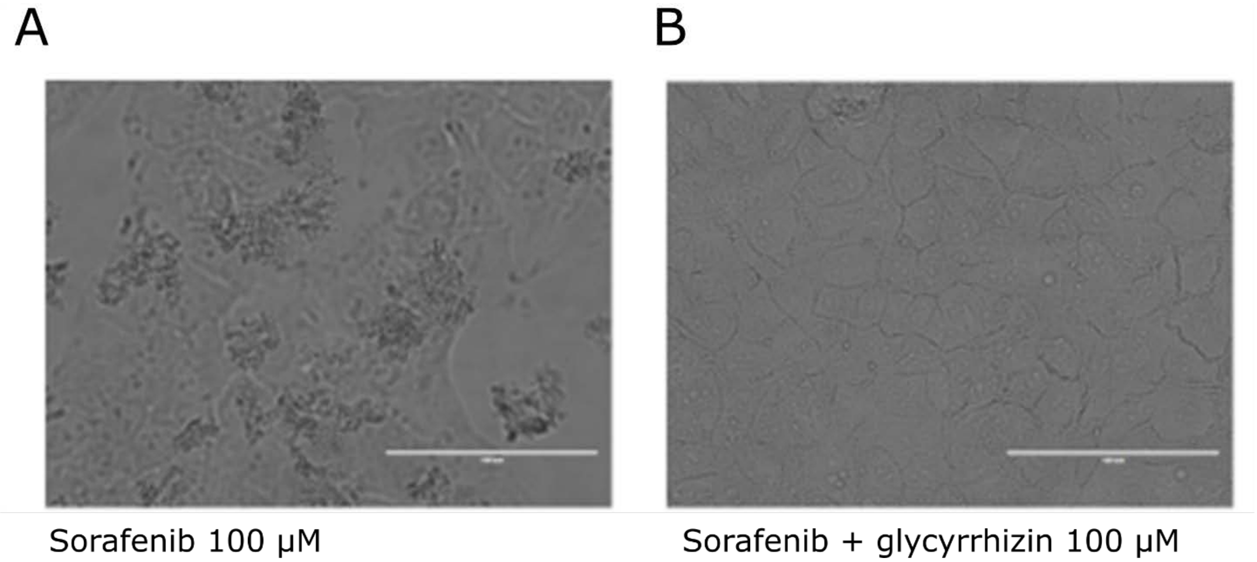


**Supplementary Figure 4:** Bright field microscopy images of HUH7 cancer cells with 100 µM sorafenib (A) vs. sorafenib-glycyrrhizin (B). Scale bar corresponds to 100 µm.

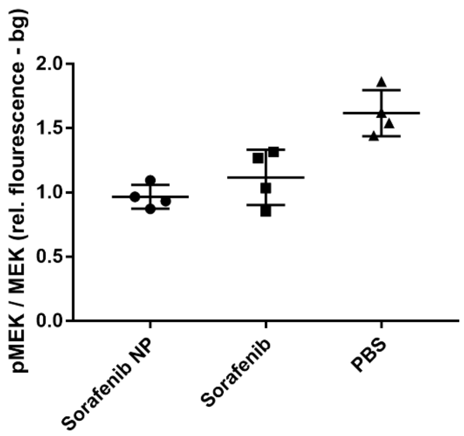


**Supplementary Figure 5**: Inhibition of MEK phosphorylation of our novel sorafenib nanoparticles is equivalent to inhibition by free sorafenib and reduces the relative amount of pMEK by 50% after 5h of treatment of HUH7 hepatocarcinoma cells. $p = 0.001$ one-way ANOVA, $p < 0.006$ Tukey's multiple comparisons test for sorafenib NP or sorafenib vs. PBS, difference between sorafenib NP and sorafenib is not significant ($p = 0.45$ Tukey's multiple comparisons test).

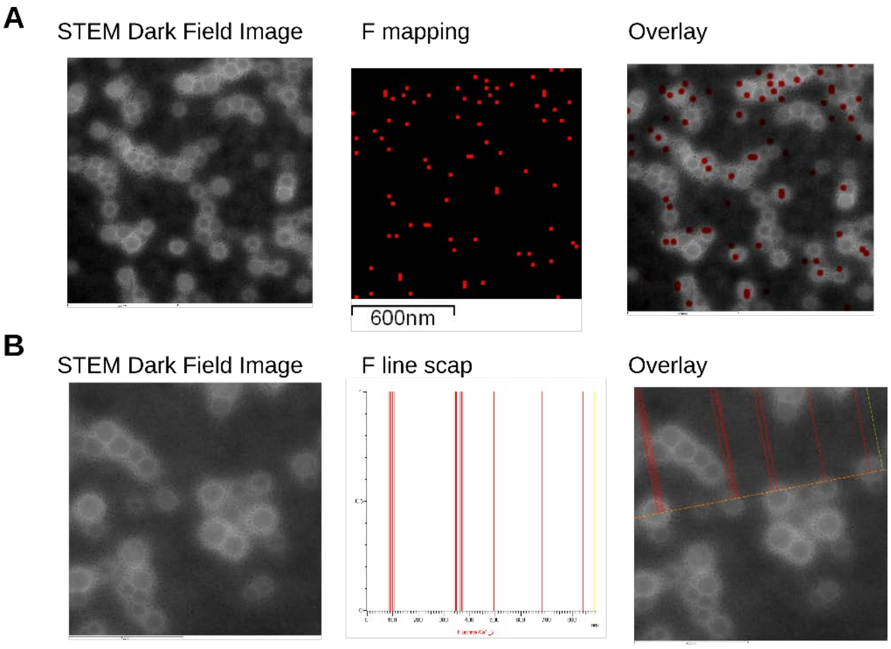

**Supplementary Figure 6:** STEM-EDS shows that sorafenib- glycyrrhizin nanoparticles are rich in fluorine from sorafenib. **A** F mapping analysis to visualize 2D distribution of fluorine in particles. **B** Line scan highlights specific enrichment of drug in the particles and lack of drug in the surrounding sample.

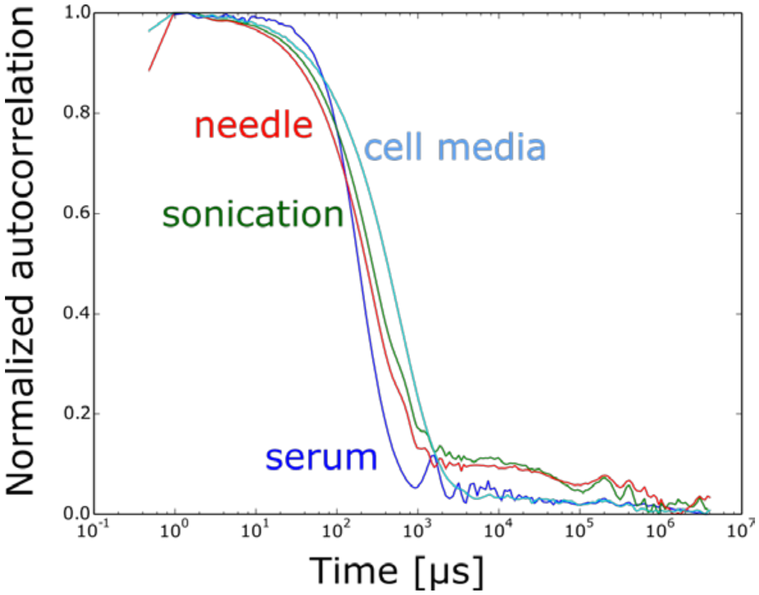


**Supplementary Figure 7**: Stability of sorafenib nanoparticles after different perturbations or in different environments. Shown are DLS autocorrelation functions of sorafenib-glycyrrhizin nanoparticles in serum (blue) or cell media (light blue) or after centrifugation and subsequent sonication for re-dispersion (green) or after pushing the nanoparticles through a 29 Gauge needle (red).

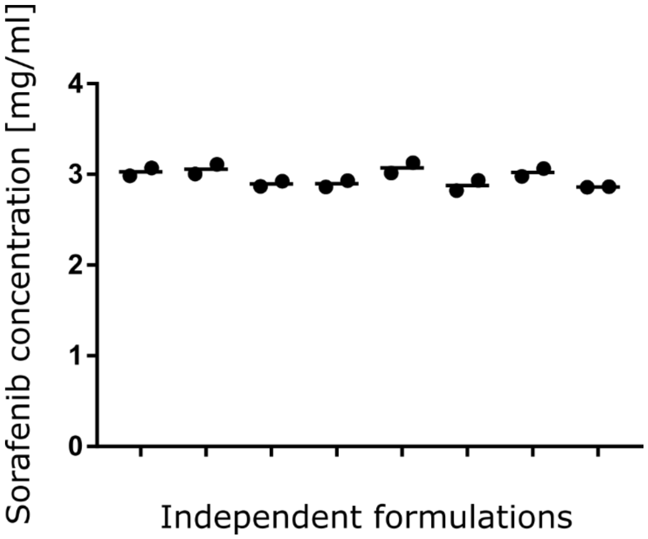


**Supplementary Figure 8**: Consistency of formulation strategy. HPLC analytics was performed in duplicates on eight independent nanoparticle formulations created to assess consistency of formulation protocol.

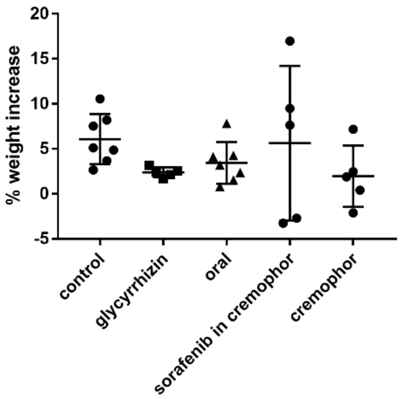


**Supplementary Figure 9:** Percentage weight increase due to tumor burden in control treatments. No significant difference was observed. One-way ANOVA $p$=0.36, $p$>0.29 Dunnett's multiple comparisons test for all treatments compared to 1% DMSO in PBS treatment control.

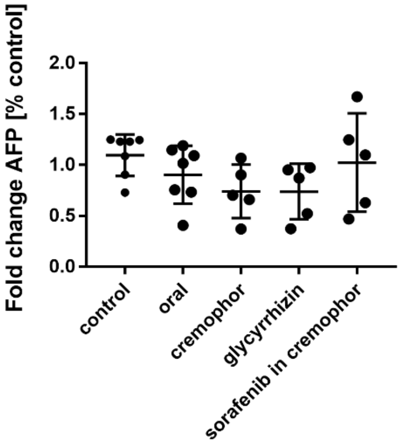


**Supplementary Figure 10:** AFP levels of control treatments. No significant difference was observed. One-way ANOVA $p$=0.209, $p$>0.17 Dunnett's multiple comparisons test for all treatments compared to 1% DMSO in PBS treatment control.

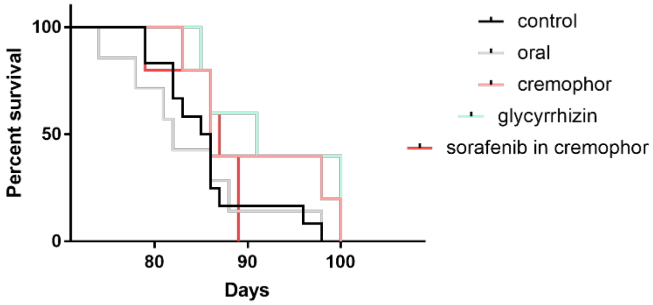


**Supplementary Figure 11**: Survival curves for control treatments, no significant difference was found. $p > 0.05$ Log-rank (Mantel-Cox) test for all treatments compared to 1% DMSO in PBS treatment control.

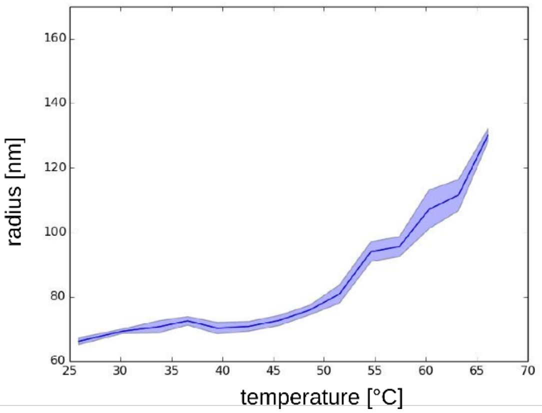


**Supplementary Figure 12:** Temperature-dependent growth of sorafenib-indomethacin particles.
